# Diverse H_2_O_2_-scavenging phenotypes of coral-associated bacteria reveal candidates for bleaching mitigation

**DOI:** 10.64898/2026.09.14.751348

**Authors:** Luanny Martins Fernandes, Talisa Doering, Wing Yan Chan, Madeleine JH van Oppen

## Abstract

There is growing interest in manipulating coral-associated bacteria to enhance coral thermal bleaching tolerance. Reactive oxygen species (ROS), particularly hydrogen peroxide (H_2_O_2_), are major drivers of coral bleaching, therefore, coral-associated bacteria capable of scavenging H_2_O_2_ may help reduce bleaching. Bacterial candidates are commonly selected through genome inference and occasionally phenotypic assays, but how well these approaches predict H_2_O_2_-scavenging capacity is unknown. We screened 29 bacterial strains isolated from the scleractinian corals *Acropora loripes* and *Galaxea fascicularis* for genome-inferred H_2_O_2_-scavenging pathways and *in vitro* activity using an ABTS radical-scavenging assay, H_2_O_2_ tolerance (the highest H_2_O_2_ concentration that allows bacterial growth), H_2_O_2_ removal from the medium, and qualitative catalase activity. Although all strains encoded at least two complete H_2_O_2_-scavenging pathways, genomic prediction did not consistently align with *in vitro* scavenging activity. Growth under H_2_O_2_ exposure was strongly coupled with H_2_O_2_ scavenging, but these traits showed no correlation with ABTS inhibition or catalase activity inferred from bubble formation. The top-ranked phenotypes, based on H_2_O_2_ tolerance and scavenging, were observed in *Roseovarius*, *Roseibium*, *Ruegeria*, and *Muricauda* strains, which tolerated 5–50 mM H_2_O_2_. These genera are common associates of microalgae, including coral endosymbionts, which are key sources of excess ROS during thermal stress. Among these, *Roseovarius* and *Muricauda* also showed high ABTS inhibition (>97%), indicating broader antioxidant capacity that may be advantageous within the complex oxidative environment of thermally stressed corals. Strains combining strong H_2_O_2_ scavenging with broad antioxidant activity may therefore represent the most promising candidates for subsequent validation within the coral holobiont.

## Introduction

Coral reefs are among the most ecologically, economically, and culturally valuable ecosystems, but are increasingly threatened by marine heatwaves causing mass coral bleaching and mortality [1–4]. Coral bleaching is the loss of the photosynthetic dinoflagellate endosymbionts (family Symbiodiniaceae) from the coral tissues, disrupting a symbiosis that fulfils most of the host’s energetic requirements through the translocation of photosynthetically fixed carbon [5, 6]. During thermal stress, disruption of photosynthetic electron transport in Symbiodiniaceae can lead to overproduction of reactive oxygen species (ROS), including singlet oxygen (^1^O_2_), superoxide (O_2_^-^), hydrogen peroxide (H_2_O_2_), and hydroxyl radicals (•OH). These ROS can cause oxidative damage in both Symbiodiniaceae and coral host cells, contributing to cellular dysfunction and ultimately the breakdown of the symbiosis that manifests as coral bleaching [7, 8]. Among the ROS involved in coral bleaching, H_2_O_2_ is especially relevant because it is relatively stable over time and membrane-permeable, allowing it to diffuse across cellular compartments [9] and potentially move between Symbiodiniaceae, coral host cells, and bacteria. H_2_O_2_ is a central intermediate in oxidative stress, produced through superoxide dismutation and capable of generating highly damaging hydroxyl radicals [10].

In addition to Symbiodiniaceae, corals host diverse microbial communities comprising other protists, bacteria, archaea, fungi, and viruses [11], collectively forming the coral holobiont [12]. Coral-associated bacteria contribute to holobiont function through nutrient cycling, antimicrobial defence, and other processes that can influence host health and stress tolerance [13, 14]. Bacterial ROS-scavenging has received increasing attention as a potentially beneficial trait within the coral holobiont, particularly under elevated temperature when oxidative stress increases [15, 16]. Genome-based studies show that many coral-associated bacteria encode extensive ROS-detoxification genes [17, 18], suggesting they may help mitigate oxidative stress within the holobiont. However, whether functional potential translates into actual ROS-scavenging phenotypes remains unknown. Bacterial H_2_O_2_ detoxification involves several enzymatic reactions (Fig. 1). Catalases decompose H_2_O_2_ into water and oxygen, whereas peroxidase-based systems reduce H_2_O_2_ to water using cellular reductants [10, 19]. Because these systems differ in substrate specificity, peroxide concentration range, cellular localisation, and redox dependence, the presence of H_2_O_2_-scavenging genes alone may not predict bacterial H_2_O_2_-scavenging capacity.

**Figure 1.**
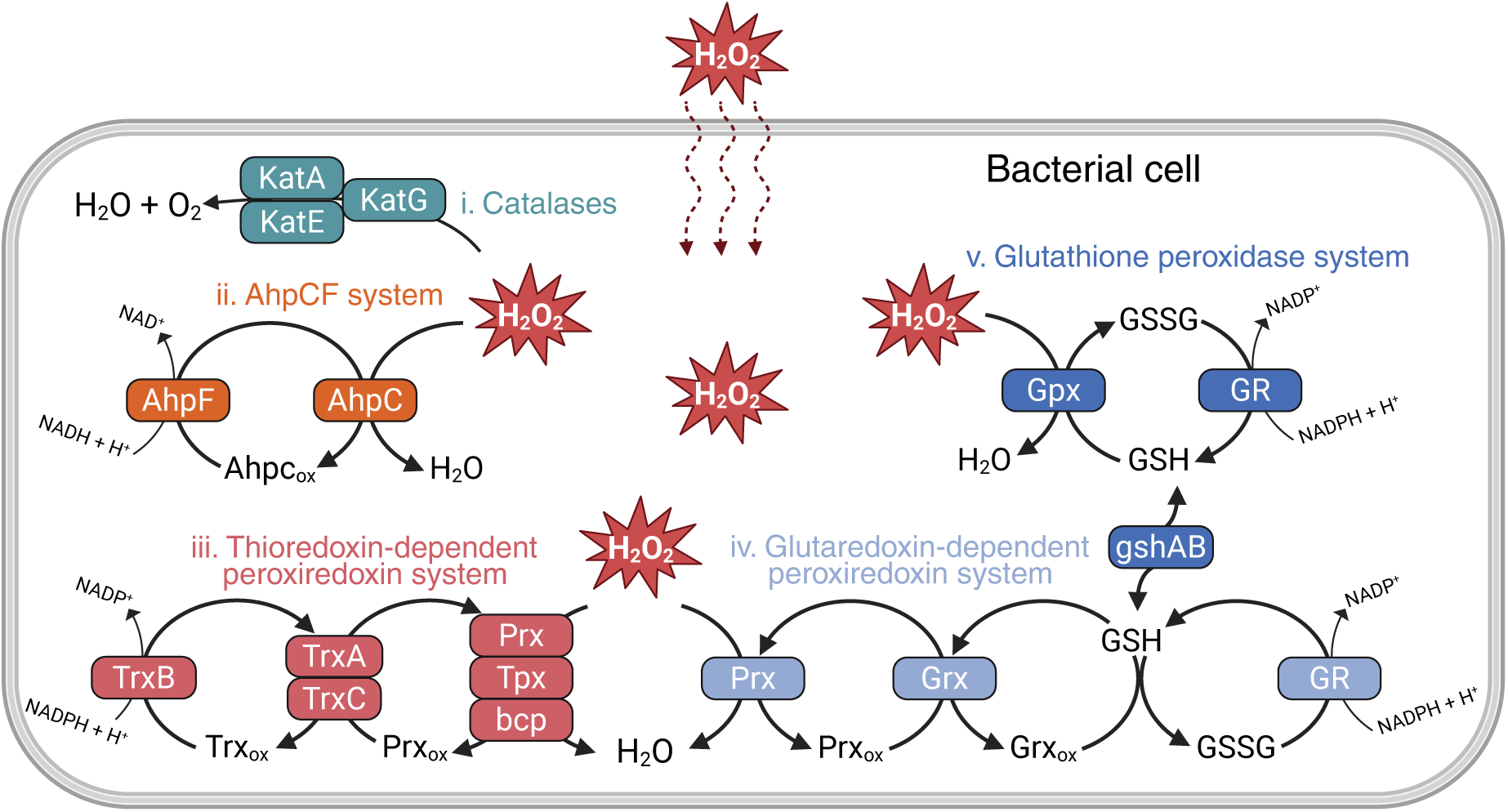
Genome-predicted H_2_O_2_-scavenging pathways across coral-associated bacterial candidates. Schematic representation of the main H_2_O_2_-scavenging mechanisms known to occur in bacterial cells. Pathways include: (i) catalases (*KatA* and *KatE*) and catalase-peroxidase (*KatG*), which convert H_2_O_2_ into H_2_O and O_2_; (ii) the alkyl hydroperoxide reductase system (AhpCF), in which *AhpC* reduces H_2_O_2_ to H_2_O and *AhpF* regenerates *AhpC* using NADH; (iii) the thioredoxin-dependent peroxiredoxin system, in which peroxiredoxin (*Prx*), thiol peroxidase (*Tpx*) and bacterioferritin comigratory protein (*bcp*) can reduce H_2_O_2_ and are regenerated by thioredoxin 1 (*TrxA*) or thioredoxin 2 (*TrxC*) through thioredoxin reductase (*TrxB*) and NADPH; (iv) the glutaredoxin-dependent peroxiredoxin system, in which *Prx* can be regenerated by glutaredoxin (*Grx*) and glutathione (GSH), which is subsequently regenerated by glutathione reductase (GR) using NADPH; and (v) the glutathione peroxidase system, in which glutathione peroxidase (*Gpx*) reduces H_2_O_2_ using GSH, producing oxidised glutathione (GSSG), which is regenerated by *GR* using NADPH. GSH biosynthesis is supported by *gshA* and *gshB*. Created in BioRender. Martins Fernandes, L. (2026) https://BioRender.com/ii0u2gg.

Manipulating coral-associated microbial communities to enhance coral thermal tolerance has emerged as a promising strategy to support coral resilience under climate change [15, 20, 21]. Several studies have shown that inoculation with putatively beneficial microorganisms can reduce bleaching or enhance survival during or following thermal stress [22–27]. However, how these bacteria confer benefits to the coral holobiont remains unclear. Current approaches for selecting candidate bacteria rely primarily on genomic prediction of function rather than assessment of relevant phenotypes [22–26, 28]. Here, we extend previous genomic characterisation and assess the *in vitro* ROS-scavenging capacity of 29 coral-associated bacterial strains isolated from *Acropora loripes* and *Galaxea fascicularis* [17, 29]. Our specific aim was to identify strains with H_2_O_2_-scavenging phenotypes that could be prioritised as candidates for mitigating oxidative stress during coral bleaching, while also determining the most reliable approaches for identifying this function. We therefore compared assays measuring broad antioxidant capacity, H_2_O_2_ tolerance, direct H_2_O_2_-scavenging, and qualitative catalase activity, with genome-inferred H_2_O_2_-scavenging potential.

## Materials and Methods

### Bacterial candidates and culture conditions

Based on previous genome-based predictions, 29 candidate strains carrying ROS-scavenging genes and pathways were selected [17, 29]. The candidate set included representatives of the genera *Endozoicomonas* (3 strains), *Neoendozoicomonas* (formerly *Endozoicomonas* Clade B; 7), *Aliiroseovarius* (1), *Muricauda* (1), *Pseudophaeobacter* (2), *Pseudovibrio* (2), *Roseibium* (1), *Roseovarius* (2), and *Ruegeria* (10) (Table 1). Culture conditions and preparation of bacterial suspensions are described in the Supplementary Methods.

**Table 1.** List of bacterial candidates.

| **Genus** | **Strain** | **Coral source** | **Reference** |
| --- | --- | --- | --- |
| *Endozoicomonas* | ALB091<br>ALB115<br>ALC013 | *Acropora loripes* | [29] |
| *Neoendozoicomonas* | ALB032<br>ALB060<br>ALB122<br>ALC020<br>ALC066<br>ALD040<br>ALD068 | *Acropora loripes* | [29] |
| *Aliiroseovarius* | DMG2200292 | *Galaxea fascicularis* | [17] |
| *Muricauda* | DMG2200308 | *Galaxea fascicularis* | [17] |
| *Pseudophaeobacter* | DMG2200302<br>DMG2200305 | *Galaxea fascicularis* | [17] |
| *Pseudovibrio* | DMG2200300<br>DMG2200345 | *Galaxea fascicularis* | [17] |
| *Roseibium* | DMG2200306 | *Galaxea fascicularis* | [17] |
| *Roseovarius* | DMG2200331<br>DMG2200361 | *Galaxea fascicularis* | [17] |
| *Ruegeria* | DMG2200288<br>DMG2200289<br>DMG2200316<br>DMG2200320<br>DMG2200324<br>DMG2200328<br>DMG2200329<br>DMG2200346<br>DMG2200347<br>DMG2200359 | *Galaxea fascicularis* | [17] |

### Re-analysis of candidate genomes for the identification of H_2_O_2_-scavenging genes and pathways

Previous studies examined broader sets of ROS-related genes and used different analytical approaches. Therefore, to specifically assess H_2_O_2_-scavenging potential across the 29 candidate strains, their published genomes [17, 29] were re-analysed. Genomes were screened for KEGG Orthology (KO) groups associated with bacterial H_2_O_2_ detoxification. Genomes were annotated using Prokka v1.14.6. KO assignments were generated from predicted proteins using KOfamScan with the KEGG KOfam HMM database, default score thresholds, and mapper output to generate gene-to-KO mappings for each genome.

A predefined list of KOs associated with bacterial H_2_O_2_-scavenging systems (Table S1) was used to generate a binary presence/absence matrix for each strain. H_2_O_2_-scavenging systems were grouped into pathway-level categories based on known bacterial H_2_O_2_-detoxification mechanisms (Fig. 1). A pathway was considered complete when all required KO components for that pathway were detected in a genome. The number of complete H_2_O_2_-scavenging pathways per strain was then calculated and used as a genome-inferred estimate of H_2_O_2_-scavenging potential.

### ABTS radical scavenging assay

Overall bacterial antioxidant capacity was assessed using an ABTS (2,2-azinobis (3-ethylbenzothiazoline-6-sulfonic acid)) radical scavenging assay modified from published protocols [30, 31]. The ABTS•+ working solution was prepared and bacterial suspensions were washed and adjusted to OD_600_ = 1.0 with 1X PBS (Supplementary Methods). Bacterial suspensions were incubated with ABTS•+ solution in 96-well plates for 30 min at 27 °C, then absorbance was measured at 734 nm. Strain-specific blanks and ABTS•+ controls were included for correction, and ABTS inhibition was calculated (Supplementary Methods).

### H_2_O_2_ tolerance assay

H_2_O_2_ tolerance was assessed by monitoring bacterial growth across seven H_2_O_2_ concentrations: 0.1, 0.5, 1, 5, 10, 20, and 50 mM. These concentrations were selected to capture a broad oxidative-stress gradient, from low H_2_O_2_ exposure (0.1 mM, previously shown to be sub-lethal in *E. coli*) to concentrations expected to strongly inhibit bacterial growth [32]. Overnight-grown bacterial cultures were diluted to OD_600_ = 0.1 using sterile-filtered Marine Broth 2216 (fMB; BD Difco^TM^, Sparks, MD, USA) and exposed to freshly prepared H_2_O_2_ in 96-well plates. Bacterial growth was monitored by measuring OD_600_ every 12 min for 60 h (CLARIOstar Plus plate reader, BMG Labtech). For each strain, maximum H_2_O_2_ tolerance was defined as the highest concentration at which growth occurred. Growth curves were blank corrected before downstream analysis.

### H_2_O_2_ depletion assay

The ability to remove H_2_O_2_ from the surrounding medium was assessed over 8 h for each strain’s maximum H_2_O_2_ concentration at which growth was previously observed. Strains that did not grow at any tested concentration were exposed to the lowest concentration of 0.1 mM H_2_O_2_. Bacterial cultures were prepared (Supplementary Methods) and diluted to OD_600_ = 0.1 using filter-sterilised reconstituted seawater (FRSW), after which they were exposed to strain-specific H_2_O_2_ concentrations. H_2_O_2_ concentration was measured immediately after H_2_O_2_ addition and after 30 min, 1 h, 1.5 h, 2 h, 4 h, and 8 h using peroxide test strips (0–100 ppm, IS124-50S; 0–400 ppm, IS123-50S; Westlab, Australia), corresponding to approximately 0–3 mM and 0–12 mM H_2_O_2_, respectively. Test-strip measurements were validated against NMR-based H_2_O_2_ quantification for two representative strains (Supplementary Methods). NMR measurements were performed according to Monakhova et al. [33]. The time to H_2_O_2_ depletion was defined as the first timepoint at which no H_2_O_2_ was detected. For strains that did not completely remove H_2_O_2_ within 8 h, the remaining concentration at the final timepoint was used to calculate the percentage removed.

### Catalase activity assay

Catalase activity was assessed indirectly using a qualitative assay based on bubble formation on the surface of the medium following Rosado et al. [22]. This assay is widely used in coral studies as a proxy for ROS-scavenging potential of bacteria [22, 23, 25, 34]. We included this assay to assess its accuracy by comparing its outcome with our H_2_O_2_ tolerance and depletion assays.

Overnight bacterial cultures (Supplementary Methods) were adjusted to OD_600_ = 1.0 using FRSW. For each strain, 3 mL of suspension was transferred to 12-well plate wells. Catalase activity was tested by adding 25 μL of 30% (v/v) H_2_O_2_ directly to each well, resulting in a final concentration of 73 mM H_2_O_2_. Strains producing visible bubbles immediately after H_2_O_2_ addition were scored as catalase-positive, and those without bubbles were catalase-negative. Unlike Rosado et al. [22], we did not assign semi-quantitative scores based on percentage of bubble coverage, as this was difficult to estimate reproducibly.

### Statistical analysis

All statistical analyses and data visualisations were performed in R version 4.5.3. (R Core Team, 2026). To test whether ABTS antioxidant capacity differed among strains, one-way analysis of variance (ANOVA) was performed, followed by Tukey’s honestly significant difference (HSD) post hoc test for pairwise comparisons.

To assess whether genome-predicted H_2_O_2_-scavenging potential correlated with phenotypic performance, Pearson correlations were calculated between the number of complete scavenging pathways and two traits: ABTS inhibition and maximum H_2_O_2_ concentration tolerated. To determine whether ABTS-measured antioxidant capacity correlated with H_2_O_2_ tolerance, Pearson correlations were estimated between ABTS inhibition and maximum H_2_O_2_ concentration tolerated. Linear regression lines with 95% confidence intervals were added to correlation plots. Maximum H_2_O_2_ tolerance was plotted on a log_10_ scale to improve visualisation across the tested range.

## Results

### Genomic potential for H_2_O_2_-scavenging across strains

Genome screening showed that, although the number and identity of pathways varied among strains, all 29 coral-associated bacterial candidates encoded at least two complete H_2_O_2_-scavenging pathways (Fig. 2). The thioredoxin-dependent peroxiredoxin pathway was complete in all strains, while other systems, including catalases, alkyl hydroperoxide reductases, glutaredoxin-dependent peroxiredoxins, and glutathione peroxidase pathways, showed strain-specific distributions (Fig. 2). The highest number of complete pathways (five) was found in all *Neoendozoicomonas* and *Pseudovibrio* strains, whereas the fewest pathways (two) occurred in *Muricauda* sp. DMG2200308 and *Ruegeria* sp. DMG2200320.

**Figure 2.**
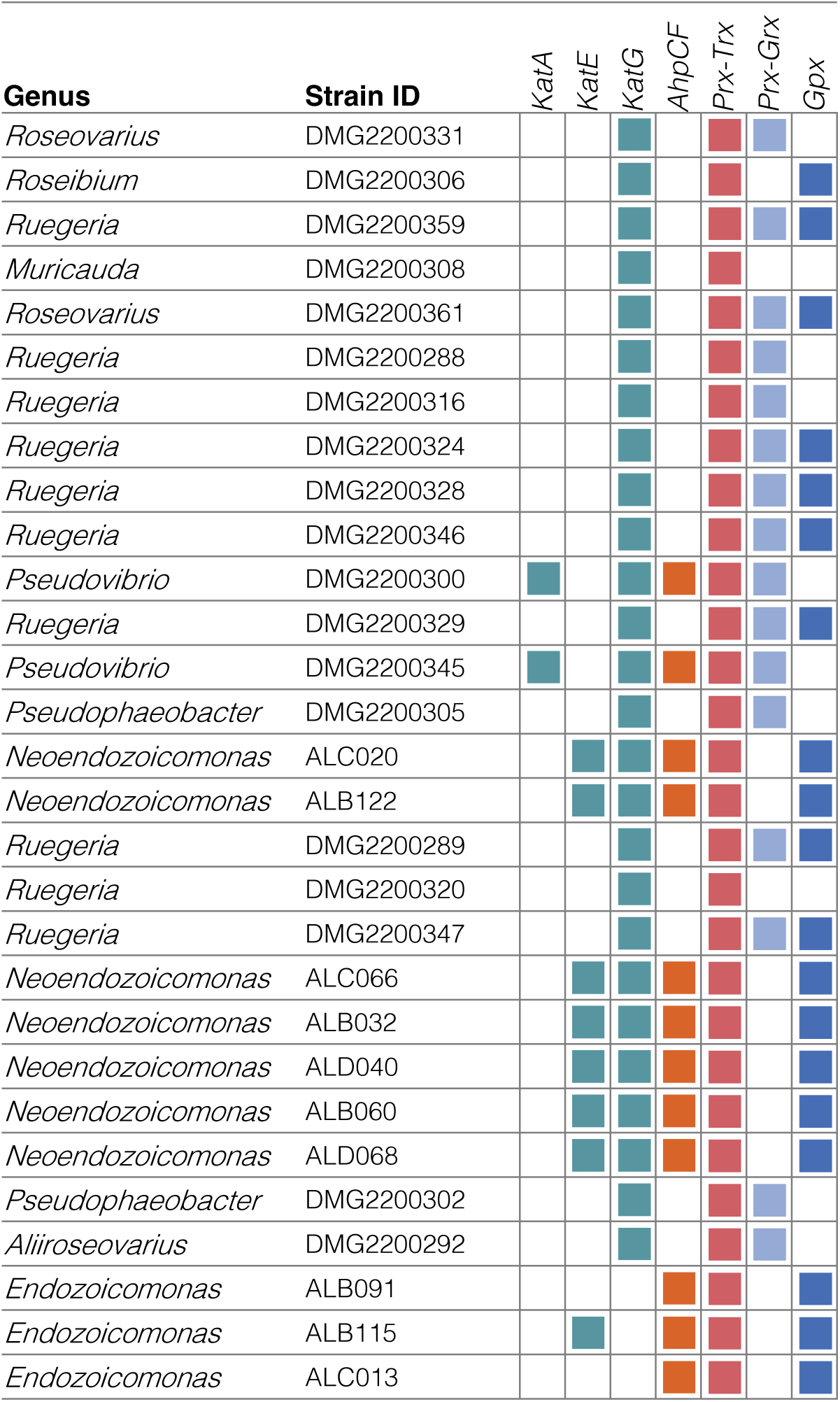
Presence of complete H_2_O_2_-scavenging pathways across the 2G candidate strains. Coloured squares indicate the presence of a complete pathway, for which all required genes were detected. The thioredoxin-dependent peroxiredoxin (*Prx-Trx*) pathway was complete in all strains, while catalase-peroxidase (*KatG*) was present in all strains except *Endozoicomonas*. Other systems showed taxon-specific distributions: *KatA* was restricted to *Pseudovibrio* strains, *KatE* to all *Neoendozoicomonas* and one *Endozoicomonas*, alkyl hydroperoxide reductase (*AhpCF*) to *Endozoicomonas, Neoendozoicomonas,* and *Pseudovibrio*. Glutathione peroxidase (*Gpx*) was detected in *Endozoicomonas*, *Neoendozoicomonas*, *Roseibium*, *Roseovarius*, and *Ruegeria*, whereas glutaredoxin-dependent peroxiredoxin (*Prx-Grx*) was present in most strains except for all *Endozoicomonas* and *Neoendozoicomonas* strains, *Roseibium* sp. DMG2200306, *Muricauda* sp. DMG2200308, and *Ruegeria* sp. DMG2200320.

### ABTS antioxidant capacity

The ABTS scavenging assay revealed substantial variation in general antioxidant capacity among the 29 bacterial candidates (Fig. 3). The highest ABTS inhibition was observed in *Aliiroseovarius* sp. DMG2200292 (100%), *Neoendozoicomonas* sp. ALC020 (98.6%), and *Roseovarius* sp. DMG2200331 (98.4%). High ABTS inhibition (>85%) was also observed in several other *Neoendozoicomonas* strains (ALD040, ALD068, ALC066), *Muricauda* sp. DMG2200308, and *Pseudovibrio* spp. DMG2200300 and DMG2200345.

**Figure 3.**
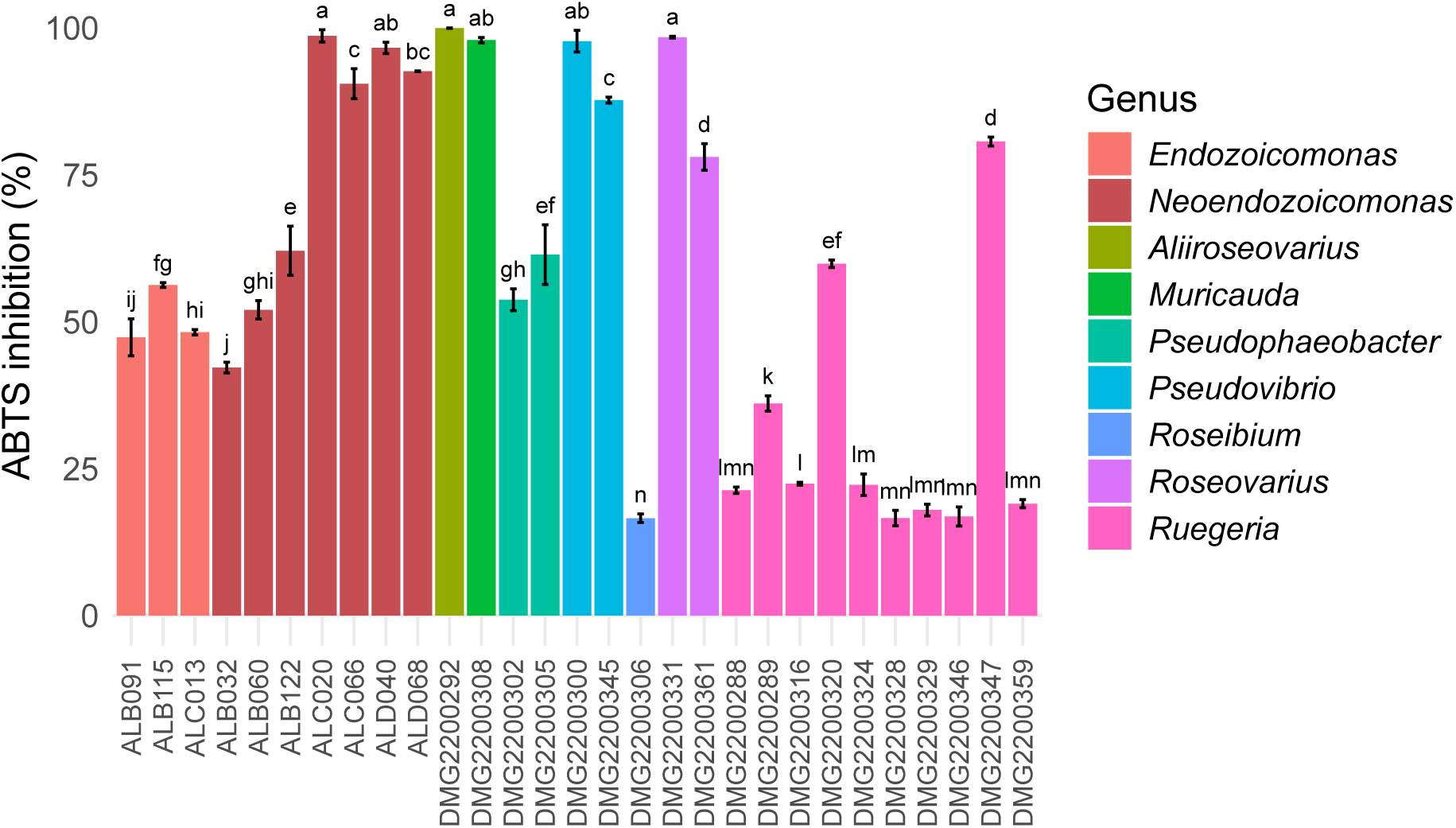
Antioxidant capacity of bacterial candidates measured by the ABTS radical scavenging assay. ABTS inhibition corresponds to the reduction in absorbance of the ABTS·+ radical relative to the control. Higher inhibition values indicate higher ABTS radical scavenging capacity. Bars show mean ± 1 SD from three biological replicates. Letters above bars indicate statistically significant differences among strains determined by one-way ANOVA followed by Tukey’s HSD test (*p* < 0.05).

In contrast, the lowest ABTS inhibition was observed in *Roseibium* sp. DMG2200306 (16.6%) and most *Ruegeria* strains (DMG2200328, DMG2200346, DMG2200329, DMG2200359, DMG2200288, DMG2200324, and DMG2200316; 16.6–22.4%).

### H_2_O_2_ tolerance and depletion

H_2_O_2_ tolerance varied markedly among strains, ranging from no detectable bacterial growth at the lowest concentration tested, 0.1 mM, to growth at up to 50 mM H_2_O_2_ (Table 2; growth curves are shown in Figs. S2 and S3). *Roseovarius* sp. DMG2200331 showed the highest tolerance, growing at 50 mM H_2_O_2_, followed by *Roseibium* sp. DMG2200306 (10 mM), and *Ruegeria* sp. DMG2200359 and *Muricauda* sp. DMG2200308 (5 mM). Twelve strains tolerated 1 mM H_2_O_2_, six 0.5 mM, four only 0.1 mM, and three strains (all *Endozoicomonas*) did not grow at any H_2_O_2_ concentration tested.

**Table 2.**
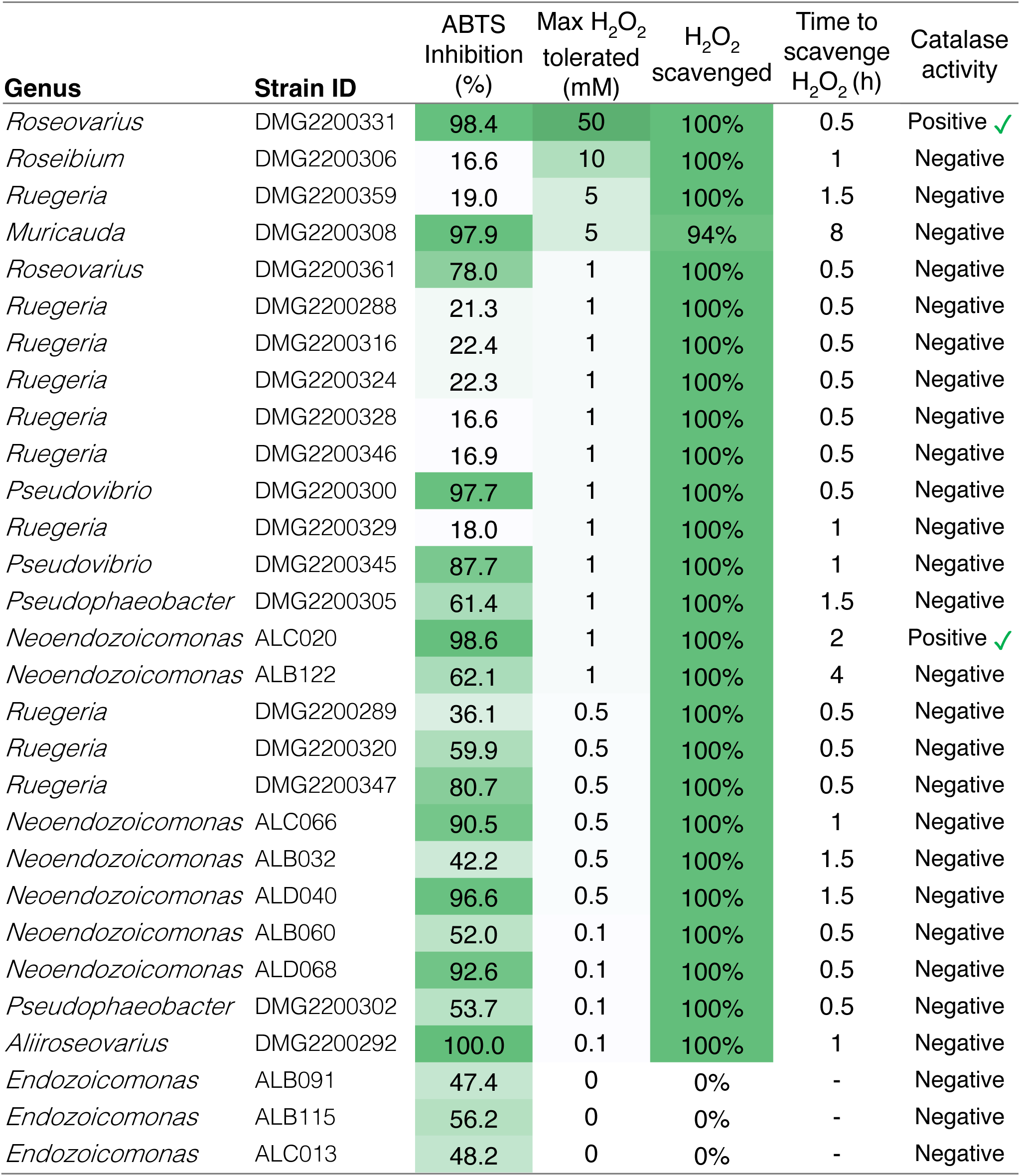
Summary of ROS-scavenging phenotypes across the 2G bacterial candidates. Columns show ABTS inhibition (%), maximum H_2_O_2_ concentration tolerated (mM), percentage of H_2_O_2_ scavenged, time to H_2_O_2_ depletion (h), and catalase activity. Green shading represents a heatmap of the values. Strains are ordered primarily by the maximum H_2_O_2_ concentration tolerated, and secondarily by the time required to scavenge H_2_O_2_, as these were used as the main criteria to identify the most robust H_2_O_2_-scavenging phenotypes.

H_2_O_2_ depletion was assessed at each strain’s maximum tolerated H_2_O_2_ concentration. All strains that grew in H_2_O_2_-supplemented medium reduced their respective H_2_O_2_ concentration within 8 h (Table 2). All strains completely depleted H_2_O_2_, most within 30 min, except *Muricauda* sp. DMG2200308, which removed approximately 94% of the initial 5 mM H_2_O_2_ within 8 h. For the three *Endozoicomonas* strains that did not grow at any H_2_O_2_ concentration, no reduction of the 0.1 mM H_2_O_2_ was observed after 8 h.

### Catalase activity

Based on the qualitative bubble assay, only *Roseovarius* sp. DMG2200331 and *Neoendozoicomonas* sp. ALC020 were catalase-positive, while all other strains were scored as catalase negative (Table 2). This included strains that showed considerable H_2_O_2_ tolerance and rapid H_2_O_2_ depletion, such as *Roseibium* sp. DMG2200306, *Ruegeria* sp. DMG2200359 and *Muricauda* sp. DMG2200308.

### Correlation analyses between predicted H_2_O_2_-scavenging potential and phenotypic traits

The number of predicted complete H_2_O_2_-scavenging pathways was not significantly correlated with either ABTS inhibition (Pearson’s *r* = 0.20, *p* = 0.29) or maximum H_2_O_2_ tolerance across all bacterial candidates (Pearson’s *r* = -0.23, *p* = 0.23; Fig. 4A–B). ABTS inhibition was also not significantly correlated with maximum H_2_O_2_ tolerance (*r* = 0.19, *p* = 0.33; Fig. 4C), and some strains showed divergent ABTS and H_2_O_2_ tolerance phenotypes. For example, *Aliiroseovarius* sp. DMG2200292 and *Neoendozoicomonas* sp. ALD040 showed high ABTS inhibition (100% and 97%, respectively) but tolerated only 0.1 and 0.5 mM H_2_O_2_, whereas *Roseibium* sp. DMG2200306 and *Ruegeria* sp. DMG2200359 showed much lower ABTS activity (17% and 19%, respectively) but tolerated higher H_2_O_2_ concentrations of 10 and 5 mM.

**Figure 4.**
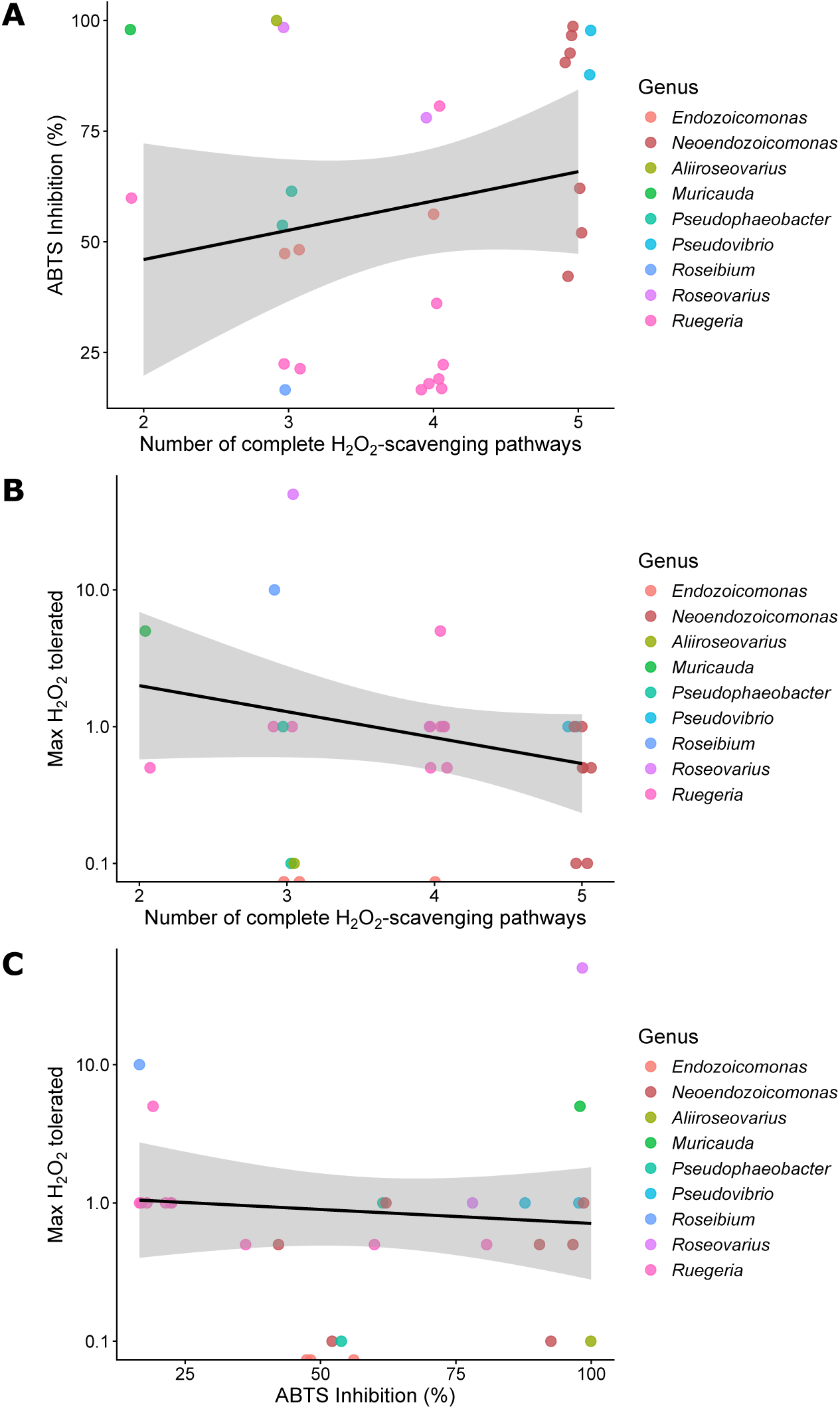
Correlation between genome-predicted H_2_O_2_-scavenging potential, ABTS antioxidant capacity and H_2_O_2_ tolerance. (A) Number of complete H_2_O_2_-scavenging pathways versus ABTS inhibition. (B) Number of complete H_2_O_2_-scavenging pathways versus maximum H_2_O_2_ tolerated. (C) ABTS inhibition versus maximum H_2_O_2_ tolerated. Lines represent linear regressions with 95% confidence intervals. Maximum H_2_O_2_ tolerance was plotted on a log_10_ scale. Pearson correlation tests showed no significant association between predicted pathway number and ABTS inhibition (*r* = 0.20, *p* = 0.29) or maximum H_2_O_2_ tolerance (*r* = -0.23, *p* = 0.23). ABTS inhibition was also not significantly correlated with maximum H_2_O_2_ tolerance (*r* = 0.19, *p* = 0.33).

## Discussion

To identify coral-associated bacteria with high-ranking H_2_O_2_-scavenging phenotypes that represent candidates for mitigating oxidative stress during coral bleaching, we compared commonly used screening approaches with direct measurements of H_2_O_2_ tolerance and depletion. This revealed several candidates: *Roseovarius* sp. DMG2200331, *Roseibium* sp. DMG2200306, *Ruegeria* sp. DMG2200359, and *Muricauda* sp. DMG2200308, which grew under high H_2_O_2_ concentrations and efficiently scavenged H_2_O_2_ from the medium. By removing H_2_O_2_, such bacteria may limit ROS accumulation within the coral holobiont, mitigating oxidative stress during thermal stress and thereby reducing bleaching. Of these, *Roseovarius* sp. DMG2200331 and *Muricauda* sp. DMG2200308 also had a high capacity to neutralise a broader spectrum of ROS, as indicated by a high level of ABTS inhibition.

### H_2_O_2_ tolerance is strongly correlated with rapid H_2_O_2_ scavenging capacity

Growth under H_2_O_2_ exposure was strongly associated with active H_2_O_2_ removal from the medium, whereas non-growing strains showed no detectable scavenging. This is relevant because a coral-associated bacterial strain may support host bleaching tolerance only if it both withstands oxidative stress and removes H_2_O_2_ from its external environment.

Since H_2_O_2_ readily crosses membranes, sustained growth in H_2_O_2_-supplemented medium likely requires defence systems that maintain intracellular H_2_O_2_ below toxic levels [10, 19, 35], making growth under H_2_O_2_ a meaningful indicator of these defences. Indeed, bacteria can withstand H_2_O_2_ stress through several mechanisms, including (1) intracellular H_2_O_2_ scavenging by catalases or peroxidases (Fig. 1), (2) neutralisation outside the cytoplasm, for example in the periplasm, thereby lowering the concentration surrounding the cell, (3) via repair of H_2_O_2_-mediated damage to proteins, Fe-S enzymes, and DNA, and (4) by limiting oxidative damage through protective metal-homeostasis responses that reduce the Fenton reaction, such as decreasing intracellular free-iron pools, and in some bacteria, replacing iron with manganese [19, 35–37].

We therefore determined whether H_2_O_2_-tolerant strains also reduced H_2_O_2_ concentrations in the medium. Because H_2_O_2_ can diffuse into bacterial cells, measured depletion may reflect intracellular detoxification, as well as periplasmic or released antioxidant enzymes and metabolites. In this study, measuring depletion did not change the ranking of candidate strains based on H_2_O_2_ tolerance, but confirmed that H_2_O_2_ tolerance was closely linked to active H_2_O_2_ removal from the medium rather than survival through intracellular repair or protective mechanisms alone. It also distinguished rapid from slower H_2_O_2_-scavengers. Therefore, H_2_O_2_ tolerance provides a practical first screen for identifying H_2_O_2_-scavenging candidates, with the depletion assay as a confirmatory step.

### ABTS and H_2_O_2_ scavenging capacity reflect different traits

ABTS neutralisation did not significantly correlate with H_2_O_2_-scavenging phenotypes. This likely reflects the nature of the assay, which measures the reduction of the pre-formed synthetic radical cation ABTS•+ and therefore broad electron- or hydrogen-donating capacity, rather than H_2_O_2_ detoxification specifically [30]. Thus, ABTS is best interpreted as a measure of general antioxidant activity, as previously used in environmental and bacterial studies [31, 38–41], rather than as a direct proxy for scavenging a specific ROS such as H_2_O_2_. ABTS•+ can be reduced by a wide range of antioxidants, many of which may not substantially contribute to H_2_O_2_ detoxification. Assay kinetics may further influence this, as chemically distinct antioxidants can reduce ABTS•+ at different rates [42]. The ABTS assay measures endpoint antioxidant capacity after a fixed incubation period, whereas the H_2_O_2_ depletion assay monitored H_2_O_2_ concentrations over a period of 8 h, providing a direct temporal assessment of bacterial H_2_O_2_ reduction. Consequently, divergent phenotypes among strains may reflect differences in the antioxidant systems each assay captured. High ABTS inhibition in strains with low H_2_O_2_ tolerance (e.g., *Aliiroseovarius* sp. DMG2200292, *Neoendozoicomonas* sp. ALD040), may indicate redox-active compounds that efficiently reduce ABTS•+ but provide insufficient H_2_O_2_ protection. Conversely, strains with lower ABTS inhibition but higher H_2_O_2_ scavenging may rely on specific H_2_O_2_-detoxifying systems not captured by the ABTS assay.

Although ABTS and H_2_O_2_ scavenging did not show a strong correlation, *Roseovarius* sp. DMG2200331 and *Muricauda* sp. DMG2200308 showed high capacity for both. A broader redox activity within the coral holobiont may be beneficial during thermal stress, because coral bleaching involves a complex oxidative environment where multiple ROS and redox-active metabolites may interact. Therefore, strains with both high H_2_O_2_ scavenging capacity and high ABTS activity may represent the most promising candidates for coral bleaching mitigation.

### Catalase activity inferred from the bubble formation assay is not a reliable method to assess bacterial H_2_O_2_ scavenging

Catalases are considered key enzymes in bacterial oxidative stress responses, decomposing H_2_O_2_ into water and oxygen. Under some assay conditions, oxygen release is visible as bubble formation after H_2_O_2_ addition, and this qualitative response, where stronger bubbling is interpreted as greater overall ROS-scavenging potential, has been widely used to select coral probiotic bacterial candidates [22, 23, 25, 34]. However, the absence of bubbles does not necessarily indicate absence of catalase activity; weak or slow activity may still produce O_2_, but too little for visible bubble formation.

Our results caution against using this assay as a proxy for overall ROS-scavenging or H_2_O_2_-scavenging specifically. Despite widespread and rapid H_2_O_2_ removal in the depletion assay, only two strains tested positive in the catalase bubble assay. Moreover, during the H_2_O_2_ depletion assay, in which each strain was challenged with the highest H_2_O_2_ concentration at which growth was still observed, no bubbles were seen immediately after H_2_O_2_ was added to the cultures. Thus, catalase activity inferred from bubble formation does not accurately reflect the H_2_O_2_-scavenging capacity of many strains.

One likely explanation is that bacterial H_2_O_2_ detoxification is not mediated by catalase alone but also by several peroxidase systems that reduce H_2_O_2_ to water using intracellular reductants (Fig. 1). This is consistent with our genomic results, which show the presence of several peroxidase-based pathways alongside catalases in the candidate strains (Fig. 2). In many bacteria, such peroxidases are the primary scavengers at low-micromolar H_2_O_2_ [19, 43, 44], which is more representative of natural levels. Notably, catalase-deficient mutants frequently resemble wild-type cells at lower H_2_O_2_, with defects emerging mainly under millimolar H_2_O_2_ challenge [44, 45]. This is relevant because H_2_O_2_ can be harmful at low-micromolar concentrations (∼10 μM) [46], and even sub-micromolar intracellular H_2_O_2_ can damage sensitive enzymes [19]. In marine environments, H_2_O_2_ typically occurs in the nanomolar range, with local increases driven by photochemistry and other biological activity [47, 48]. At coral surfaces under ambient conditions, external H_2_O_2_ varies among taxa: ∼500 nM for *Porites*, ∼250 nM for *Pocillopora*, and near zero for *Fungia* [48]. While corals regulate nanomolar H_2_O_2_ concentrations under non-stress conditions, H_2_O_2_ levels at coral surfaces and within tissues during thermal stress remain unknown.

This mismatch between ecologically relevant H_2_O_2_ concentrations and the dose used in the catalase bubble assay raises additional methodological concern. The assay relies on acute exposure to high H_2_O_2_ concentrations (approximately 70–90 mM), far above the tolerance range of all strains tested. Because H_2_O_2_ tolerance varied substantially, sensitive strains may fail to produce bubbles even if they encode catalases or can degrade H_2_O_2_ at lower, more physiologically relevant levels. In this sense, the assay may underestimate catalase activity and generate false negatives, including through enzyme inactivation or cellular damage [19, 49].

Other studies have attempted to improve catalase detection by adding Triton X-100, which stabilises the catalase-derived oxygen as foam and allows semi-quantitative scoring from foam height [50]. Although this may improve visual detection of catalase-associated oxygen production, it does not overcome the broader limitation of using a single-enzyme, high-dose assay to infer overall H_2_O_2_ detoxification capacity.

### Genomic prediction poorly reflects *in vitro* phenotypes

Genome-based screening has been used as an attractive, cost- and time-efficient way to identify candidate bacteria for coral microbiome manipulation and to predict functions linked to oxidative stress alleviation, nutrient provision, colonisation and persistence in the coral holobiont [15, 17, 29, 51].

In our study, genome-predicted potential did not reliably reflect ROS-scavenging performance *in vitro*. Despite all strains encoding complete ROS-scavenging pathways, phenotypic assays revealed substantial variation in overall antioxidant capacity, H_2_O_2_ tolerance and scavenging, and catalase activity. No single genomic pathway consistently distinguished high-from low-performing strains, and the number of complete H_2_O_2_-scavenging pathways did not predict H_2_O_2_ tolerance or scavenging (Fig. 4). This mismatch is biologically plausible since effective H_2_O_2_ detoxification depends not simply on gene presence, but also on context-dependent gene expression, enzyme activity, reductant supply and overall physiological capacity [19, 35]. H_2_O_2_-scavenging can be mediated by multiple overlapping systems that differ in substrate range, cellular localisation, and effective concentration range [10, 19, 52].

Whether *in vitro* and *in hospite* phenotypes are similar is currently unknown. This critical knowledge gap could be addressed by examining H_2_O_2_-scavenging phenotypes of genomically characterised bacteria *in hospite* or at a minimum in the presence of the coral holobiont metabolites or exudates [53]. The former is challenging but could perhaps be obtained via metatranscriptomic approaches. The latter could be explored via the use of co-culture devices that permit bacterial exposure to holobiont exudates.

### Phenotypic screening identified usual suspects as promising candidates

The strains with the highest H_2_O_2_-scavenging capacity identified here belong to *Roseovarius*, *Roseibium*, *Ruegeria*, and *Muricauda*. These genera are frequently linked to microalgae- or Symbiodiniaceae-associated environments, where they can support algal growth, photophysiology, nutrient exchange and stress tolerance. For example, *Roseovarius* inoculation improved thermal tolerance of the coral photosymbiont *Breviolum minutum*, potentially through ROS scavenging [54]. *Roseibium* (formerly *Labrenzia*) engaged in metabolic exchanges with Symbiodiniaceae and enhanced their growth [55]. Both *Roseibium* and *Muricauda* were linked to improved Symbiodiniaceae photophysiology under acute temperature and light stress [56]. *Muricauda* also reduced ROS and restored photosynthetic performance in thermally stressed Symbiodiniaceae cultures, likely through zeaxanthin production [57]. *Ruegeria* has been shown to support Symbiodiniaceae nutrition through nitrogen transfer [58].

The association of these genera with Symbiodiniaceae is highly relevant for coral bleaching, because Symbiodiniaceae are a major site of ROS production during thermal stress [7]. Bacteria that closely associate with Symbiodiniaceae or their phycosphere [59] may be well positioned to remove H_2_O_2_ near its site of production, limiting local accumulation or diffusion into coral tissues. Symbiodiniaceae-associated bacteria with strong H_2_O_2_-scavenging phenotypes therefore could help reduce oxidative stress at the host-symbiont interface and contribute to holobiont resilience during thermal stress.

*Roseovarius* may be a strong candidate because, in addition to enhancing Symbiodiniaceae thermal tolerance [54], members of this genus have been linked to ROS-scavenging traits and sulphur metabolism. *Roseovarius* participates in dimethylsulfoniopropionate (DMSP) metabolism and dimethylsulfide (DMS) production in algal-associated systems, including interactions with the dinoflagellate *Pfiesteria piscicida* [60, 61]. DMS and DMSP are important components of the cell’s antioxidant system [62].

*Roseibium* is a recurrent and functionally important member of Symbiodiniaceae-associated microbiomes. It is a core bacterial genus across cultured *Symbiodinium* strains, accounting for up to 38.4% of their bacterial community [63], and is consistently associated with other cultured genera of Symbiodiniaceae [64]. *Roseibium* has been detected in intracellular bacterial communities of Symbiodiniaceae freshly isolated from *G. fascicularis* [65, 66], and was reported to increase in relative abundance in Symbiodiniaceae cultures under heat stress [67]. Previous work identified a *Roseibium* strain isolated from the anemone *Exaiptasia diaphana* as a high free-radical-scavenging isolate [34]. Co-culture experiments showed reciprocal growth benefits between *Roseibium* and Symbiodiniaceae under vitamin B_12_ limitation [65]. Finally, members of this genus participate in DMSP biosynthesis and degradation [68], suggesting an additional potential pathway to influence redox processes in Symbiodiniaceae.

*Muricauda* is also detected across multiple Symbiodiniaceae cultures [63, 64]. Members of this genus are known for their carotenoid production, including zeaxanthin, which has antioxidant and photoprotective properties [57]. *Muricauda* strains isolated from *G. fascicularis* and surrounding seawater encode complete zeaxanthin biosynthesis genes and produce several carotenoids [69]. Notably, *Muricauda* sp. DMG2200308, tested here, was previously identified as a probiotic candidate encoding zeaxanthin and β-carotene biosynthesis genes [17].

In contrast to *Roseibium* and *Muricauda*, *Ruegeria* is less commonly reported in cultured Symbiodiniaceae microbiomes, but several studies support its relevance as an algal- and coral-associated bacterium with potentially beneficial functions. In algal systems, *Ruegeria pomeroyi* is known to support growth of the diatom *Thalassosira pseudonana* through vitamin B_12_ provision while metabolising algal-derived sulfonates [70]. *Ruegeria* was present in intracellular bacterial communities of *in hospite* Symbiodiniaceae isolated from *G. fascicularis* [66] and is a common coral symbiont [71] that is frequently reported to increase in relative abundance under stress conditions [72, 73], suggesting a role in coral stress responses. It has also been linked to pathogen suppression: *Ruegeria* isolates from *G. fascicularis* inhibit the coral pathogen *Vibrio coralliilyticus in vitro* [74], and *Ruegeria profundi* reduces *V. coralliilyticus*-induced bleaching and dysbiosis in corals during pathogen challenge [75]. Further, members of this genus are involved in DMSP sulphur cycling, potentially relevant in coral-associated environments [76].

Collectively, these reports suggest that the most promising H_2_O_2_-scavenging candidates identified here are not only high-performing *in vitro* but also belong to bacterial genera with plausible ecological roles supporting algal symbiosis and coral holobiont stress resilience.

### *Neoendozoicomonas*: a tissue-associated candidate genus with moderate H_2_O_2_-scavenging capacity

Tissue-associated bacteria may be especially valuable for long-term bleaching mitigation as they are more likely to form stable associations with the coral host than transient bacteria in the mucus layer [20, 21]. Although *Roseovarius*, *Roseibium*, *Ruegeria* and *Muricauda* showed strong *in vitro* H_2_O_2_-scavenging capacity and have been linked to Symbiodiniaceae-associated environments, their precise localisation within the coral holobiont remains poorly resolved. The closely related genera *Endozoicomonas* and *Neoendozoicomonas* provide a contrasting example in which tissue-localisation and persistence are well supported, but H_2_O_2_-scavenging performance ranged from low to moderate. Members of these genera are widespread and often dominant coral-associated bacteria [77, 78], frequently occurring as cell-associated microbial aggregates (CAMAs) in coral tissues [79–81]. In *Acropora loripes*, the same *Endozoicomonas* and *Neoendozoicomonas* strains tested here were shown to form CAMAs within the gastrodermis, with *Neoendozoicomonas* accounting for most CAMAs detected by clade-specific FISH [81]. Gastrodermal tissues contain Symbiodiniaceae, placing these strains close to the host-symbiont interface where oxidative stress is expected to increase during bleaching.

Thus, despite showing only moderate H_2_O_2_ tolerance in our assays, *Neoendozoicomonas* warrants further examination as a bleaching-mitigating candidate. By contrast, *Endozoicomonas* strains showed no tolerance even at the lowest H_2_O_2_ concentration tested. Although the number of complete H_2_O_2_-scavenging pathways did not predict the observed phenotypic performance across all strains, the better performance of *Neoendozoicomonas* is consistent with its higher number of pathways relative to the closely related *Endozoicomonas*, as well as broader genomic differences in predicted metabolic and host-interaction traits [29].

Members of both genera likely play additional beneficial roles within their coral hosts, including nutrient cycling, DMSP metabolism, and vitamin and amino acid provisioning [18, 29, 78–80]. Recent inoculation studies confirm the capacity of *Endozoicomonas* to colonise coral tissues, with *E. acroporae* forming CAMAs in newly settled *Acropora kenti* and in *Stylophora pistillata* following experimental inoculation [82, 83]. Thus, while the stronger H_2_O_2_-scavenging candidates may have greater immediate potential for mitigating oxidative stress during coral bleaching, *Neoendozoicomonas* may still be able to relieve some oxidative stress while providing additional beneficial functions. Its capacity to colonise coral tissues and form persistent host associations may allow these benefits to be delivered over protracted periods of time.

### Implications for identifying bacterial probiotic candidates for alleviating coral bleaching

Previous coral probiotic consortia have been selected using genomic evidence for potentially beneficial functions, phenotypic pathogen-inhibition assays, and catalase activity inferred from bubble formation as a proxy for ROS scavenging [22, 23]. More recently, Tang et al. [25] specifically selected a ROS-scavenging candidate based on genomic evidence and the catalase bubble assay, while other studies used previously selected [24, 28] or host-tailored consortia [26], rather than establishing new screening pipelines. However, our findings show that gene presence and catalase activity inferred from bubble formation do not represent reliable indicators of H_2_O_2_ detoxification, and these commonly used screening approaches may misclassify H_2_O_2_-scavenging potential of coral-associated bacteria. While ABTS inhibition was also found to inaccurately reflect H_2_O_2_ detoxification, it was likely because this assay instead quantifies a broader ROS scavenging activity. The direct phenotypic assays, H_2_O_2_ tolerance and depletion, in contrast, provide a cost- and time-efficient approach for identifying H_2_O_2_-detoxifying candidates that may mitigate coral bleaching. Candidate strains that also exhibit high ABTS inhibition are expected to have the best ability to reduce oxidative stress in corals during summer heatwaves.

Moving forward, targeted *in vitro* phenotypic screening should be followed by validation in the presence of the coral holobiont. Because traits measured in culture may not fully reflect function during interaction with the coral host and its algal symbionts, candidates should be tested under bleaching-relevant conditions to determine whether they reduce oxidative stress and improve holobiont performance. Transcriptomics could then link observed benefits to specific ROS-scavenging pathways expressed under H_2_O_2_ exposure, thermal stress, or host-associated conditions. If *in hospite* validations confirm that *in vitro* H_2_O_2_ tolerance and depletion assays predict holobiont benefits, these assays could provide a practical framework for selecting bacterial candidates to support coral resilience during bleaching events.

## Acknowledgements

We would like to thank Ivanhoe Leung and Evelyn Huang for their assistance with the NMR measurements.

## Data availability

All data generated or analysed during this study are included in this published article and its supplementary information files.

