## Supplemental Material for "Diverse H_2_O_2_-scavenging phenotypes of coral-associated bacteria reveal candidates for bleaching mitigation"

**This file includes:**

- **Supplementary Methods**
- **Table S1**
- **Figures S1 to S3**

### Supplementary Methods

#### Bacterial culture conditions

The 29 coral-associated bacterial strains had been cryopreserved by freezing cells in 40% (v/v) glycerol in liquid nitrogen ($LN_2$), followed by storage at -80 °C in the Marine Microbial Symbiont Facility at the University of Melbourne. Strains were grown on Marine Agar 2216 and in Marine Broth 2216 (BD Difco™, Sparks, MD, USA). Marine Broth was sterile-filtered through a 0.22 µm membrane before use to remove suspended particles that could interfere with absorbance-based measurements. The use of filtered Marine Broth (fMB) was validated by comparing bacterial growth in filtered and unfiltered Marine Broth, which showed comparable growth for all strains.

For all assays, single colonies were used to inoculate 10 mL of fMB. Cultures were incubated for 22 h with shaking at 200 rpm. *Endozoicomonas* and *Neoendozoicomonas* strains were incubated at 28 °C, whereas *Aliiroseovarius*, *Muricauda*, *Pseudophaeobacter*, *Pseudovibrio*, *Roseibium*, *Roseovarius*, and *Ruegeria* strains were incubated at 26 °C.

#### ABTS radical scavenging assay

The ABTS•+ working solution was prepared by mixing 7 mM ABTS with 2.45 mM potassium persulfate at a 1:1 ratio and incubating in the dark at room temperature for 14 h. Before use, the ABTS•+ solution was diluted with 1X phosphate-buffered saline (PBS) to an absorbance of approximately 0.7 at 734 nm.

Bacterial cultures grown overnight were harvested by centrifugation at 3,000 × g at 4 °C for 30 min. After removal of the supernatant, the cell pellet was washed twice by resuspension in 10 mL of 1X PBS and centrifugation at 3,000 × g at 4 °C for 15 min. Following the second wash, the pellet was resuspended in 1.5 mL of 1X PBS. The optical density of each bacterial suspension was measured at 600 nm using a CLARIOstar Plus plate reader ($OD_{600}$, BMG Labtech), and cell densities were adjusted to $OD_{600}$ = 1.0 with 1X PBS.

For each strain, 100 µL of bacterial suspension was added to a 96-well plate in triplicate, followed by 100 µL of ABTS•+ solution. Blanks containing 100 µL of bacterial suspension and 100 µL of 1X PBS were included to account for strain-specific background absorbance. Negative controls containing 100 µL of ABTS•+ solution and 100 µL of 1X PBS were included to represent maximum ABTS•+ absorbance. Plates were incubated in the dark at 27 °C for 30 min, after which absorbance was measured at 734 nm. ABTS inhibition was calculated after blank correction using the following equation:

$$ABTS\ Inhibition\ (\%) = \left(\frac{A_{control} - A_{sample}}{A_{control}}\right) x\ 100$$

where $A_{control}$ is the absorbance of the ABTS•+ negative control and $A_{sample}$ is the blank-corrected absorbance of the bacterial sample.

**$H_2O_2$ tolerance assay**

Overnight bacterial cultures were diluted to $OD_{600}$ = 0.1 in sterile-filtered Marine Broth 2216 (fMB; BD Difco™, Sparks, MD, USA). Bacterial strains were exposed to 7 $H_2O_2$ concentrations: 0.1, 0.5, 1, 5, 10, 20, and 50 mM. $H_2O_2$ solutions were prepared fresh immediately before each assay to minimise degradation before bacterial exposure.

For each strain and $H_2O_2$ concentration, cultures were loaded into 96-well plates in triplicate, after which the corresponding $H_2O_2$ solutions were added. Wells were mixed by gentle pipetting immediately after $H_2O_2$ addition to ensure even distribution before the plate was transferred to the plate reader. Wells containing each $H_2O_2$ concentration in fMB without bacterial cells were included as blanks to account for any effect of $H_2O_2$ on absorbance readings.

Bacterial growth was monitored by measuring $OD_{600}$ every 12 min for 60 h using a plate reader. Plates were incubated at 28 °C for *Endozoicomonas* and *Neoendozoicomonas* strains and at 26 °C for the remaining strains, with double-orbital shaking at 300 rpm between each reading.

**$H_2O_2$ depletion assay**

Overnight bacterial cultures were centrifuged at 3,000 × g at 4 °C for 30 min to pellet bacterial cells. Supernatants were discarded, and cells were washed twice with FRSW before being resuspended in FRSW. The $OD_{600}$ of each bacterial suspension was measured and adjusted to $OD_{600}$ = 0.1 with FRSW.

Filtered reconstituted seawater (FRSW) was prepared using reverse osmosis water and Red Sea Salt™ (Red Sea, USA) to a salinity of 35 parts per thousand (ppt). The solution was filter-sterilised through a 0.22 µm membrane filter.

$H_2O_2$ was added to each bacterial suspension at the strain-specific concentration determined from the tolerance assay. $H_2O_2$ concentration was measured immediately after $H_2O_2$ addition and subsequently after 30 min, 1 h, 1.5 h, 2 h, 4 h, and 8 h using peroxide test strips. Peroxide test strip measurements were validated against a nuclear magnetic resonance (NMR)-based $H_2O_2$ quantification for two representative strains: *Roseovarius* sp. DMG2200331, which tolerated the highest tested concentration of 50 mM $H_2O_2$, and *Endozoicomonas* sp. ALC013, which did not grow at any $H_2O_2$ concentration tested. Because $H_2O_2$ concentrations measured using test strips were consistent with the NMR-based measurements (Fig. S1), peroxide test strips were used for all strains as a more practical and time-efficient method for monitoring $H_2O_2$ concentration.

The percentage of $H_2O_2$ that was removed by each bacterial strain was calculated as:

$$H_2O_2\ removed\ (\%) = \left(\frac{C_{initial} - C_{final}}{C_{initial}}\right) x\ 100$$

where $C_{initial}$ is the strain-specific $H_2O_2$ concentration added to the bacterial suspension, and $C_{final}$ is the concentration remaining at the final measurement timepoint.

**Catalase activity assay**

For the catalase activity assay bacterial cells were prepared as described in the $H_2O_2$ depletion assay and adjusted to $OD_{600}$ = 1.0 with FRSW.

**Table S1.** KEGG Orthology annotations associated with bacterial peroxide stress response pathways.

| KO | Gene | Protein/Enzyme |
|---|---|---|
| K03781 | KatA, KatB, KatE | catalase |
| K03782 | KatG | catalase-peroxidase |
| K07217 | KatN | manganese catalase |
| K03386 | ahpC, PRDX2_4 | peroxiredoxin 2/4 |
| K24119 | ahpC | NADH-dependent peroxiredoxin subunit C |
| K24126 | ahpC | lipoyl-dependent peroxiredoxin subunit C |
| K03387 | ahpF | NADH-dependent peroxiredoxin subunit F |
| K24158 | prx | thioredoxin-dependent peroxiredoxin |
| K24157 | prxII | thioredoxin-dependent peroxiredox |
| K11065 | tpx | thioredoxin-dependent peroxiredoxin |
| K03564 | bcp, PrxQ | thioredoxin-dependent peroxiredoxin |
| K03671 | trxA | thioredoxin |
| K03672 | trxC | thioredoxin 2 |
| K00384 | trxB | thioredoxin reductase |
| K24137 | prx1 | glutaredoxin/glutathione-dependent peroxiredoxin |
| K24138 | prx3 | glutaredoxin/glutathione-dependent peroxiredoxin |
| K24140 | prxII | glutaredoxin-dependent peroxiredoxin |
| K03674 | grxA | glutaredoxin 1 |
| K03675 | grxB | glutaredoxin 2 |
| K03676 | grxC | glutaredoxin 3 |
| K07390 | grxD | monothiol glutaredoxin |
| K00432 | gpx | glutathione peroxidase |
| K05361 | gpx4 | phospholipid-hydroperoxide glutathione peroxidase |
| K00383 | GSR | glutathione reductase |
| K01919 | gshA | glutamate-cysteine ligase |
| K06048 | gshA | glutamate-cysteine ligase / carboxylate-amine ligase |
| K01920 | gshB | glutathione synthase |
| K04761 | oxyR | LysR family transcriptional regulator, hydrogen peroxide-inducible genes activator |
| K09825 | PerR | Fur family transcriptional regulator, peroxide stress response regulator |

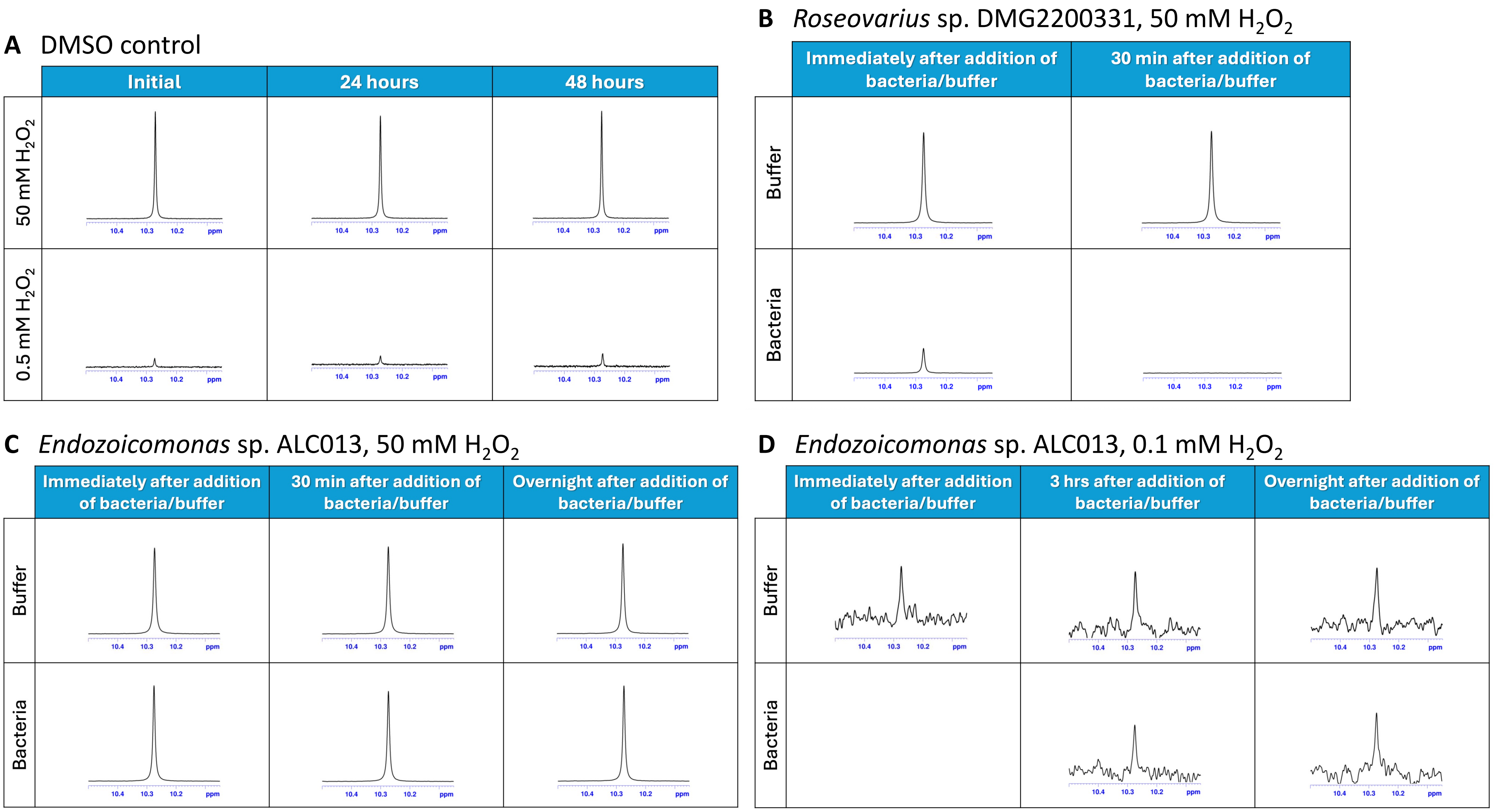


**Figure S1.** NMR validation of $H_2O_2$ detection and depletion by bacterial cells. (A) Stability of $H_2O_2$ in DMSO-$d_6$. $H_2O_2$ was mixed with DMSO-$d_6$ at either 50 mM or 0.5 mM and measured immediately, after 24 h, and after 48 h. The $H_2O_2$ peak remained stable over time, indicating minimal reaction between $H_2O_2$ and DMSO-$d_6$. (B) NMR spectra showing depletion of 50 mM $H_2O_2$ by *Roseovarius* sp. The $H_2O_2$ peak was reduced immediately after addition of bacteria and was no longer detectable after 30 min, whereas the buffer control remained unchanged. (C) NMR spectra showing that *Endozoicomonas* sp. ALC013 did not deplete 50 mM $H_2O_2$. $H_2O_2$ peak remained similar in the bacterial treatment and buffer control over time. (D) NMR spectra showing that *Endozoicomonas* sp. ALC013 did not deplete 0.1 mM $H_2O_2$. $H_2O_2$ peak remained similar in the bacterial treatment and buffer control over time.

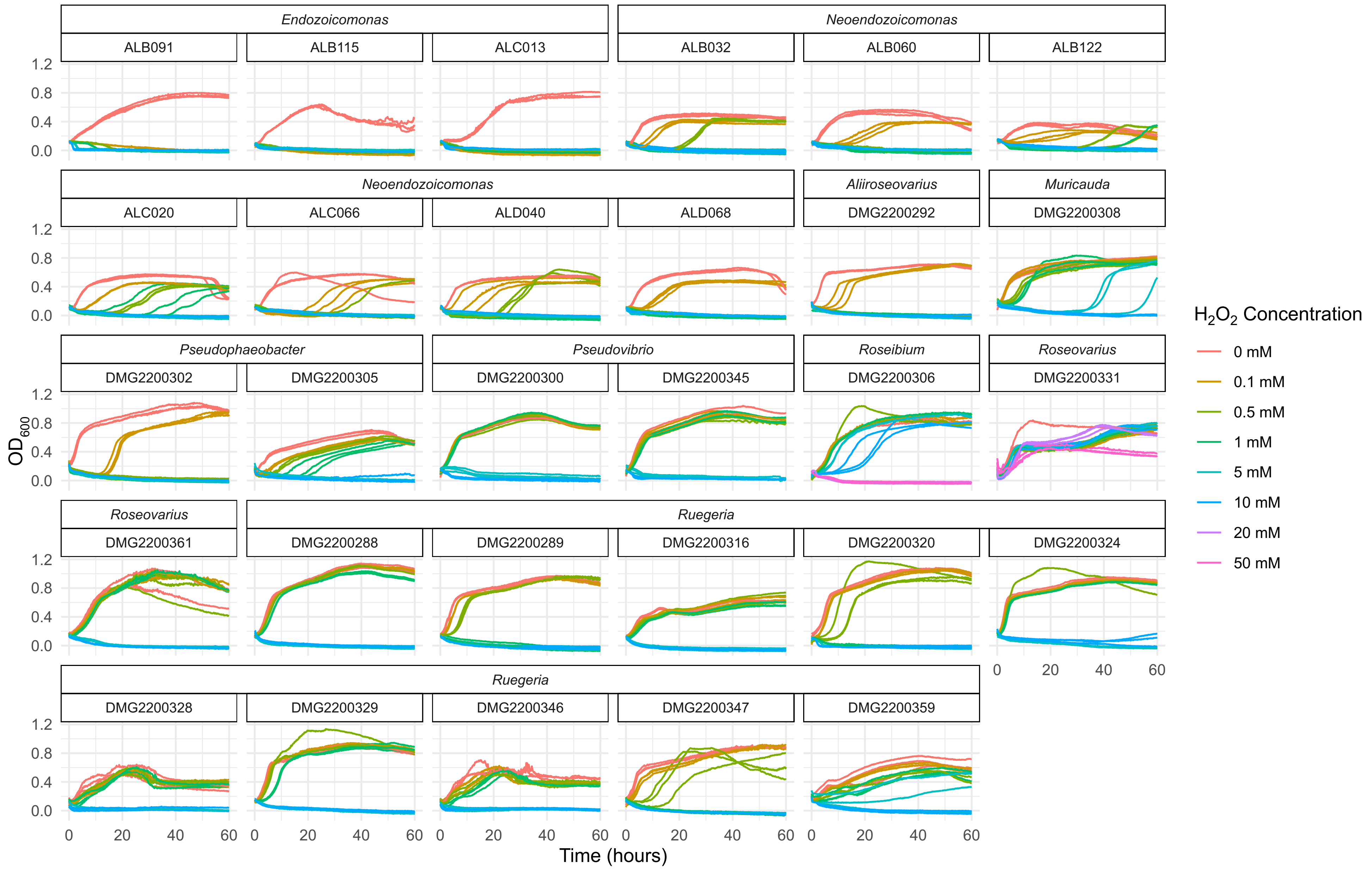


**Figure S2.** Growth curves of the 29 bacterial strains exposed to $H_2O_2$ concentrations ranging from 0 to 50 mM in 0.22 µm-filtered Marine Broth. Optical density at 600 nm ($OD_{600}$) measurements were taken every 12 minutes for 60 hours using a CLARIOstar Plus plate reader. *Endozoicomonas* and *Neoendozoicomonas* strains were incubated at 28 °C, whereas all other genera were incubated at 26 °C. Lines represent individual biological replicates.

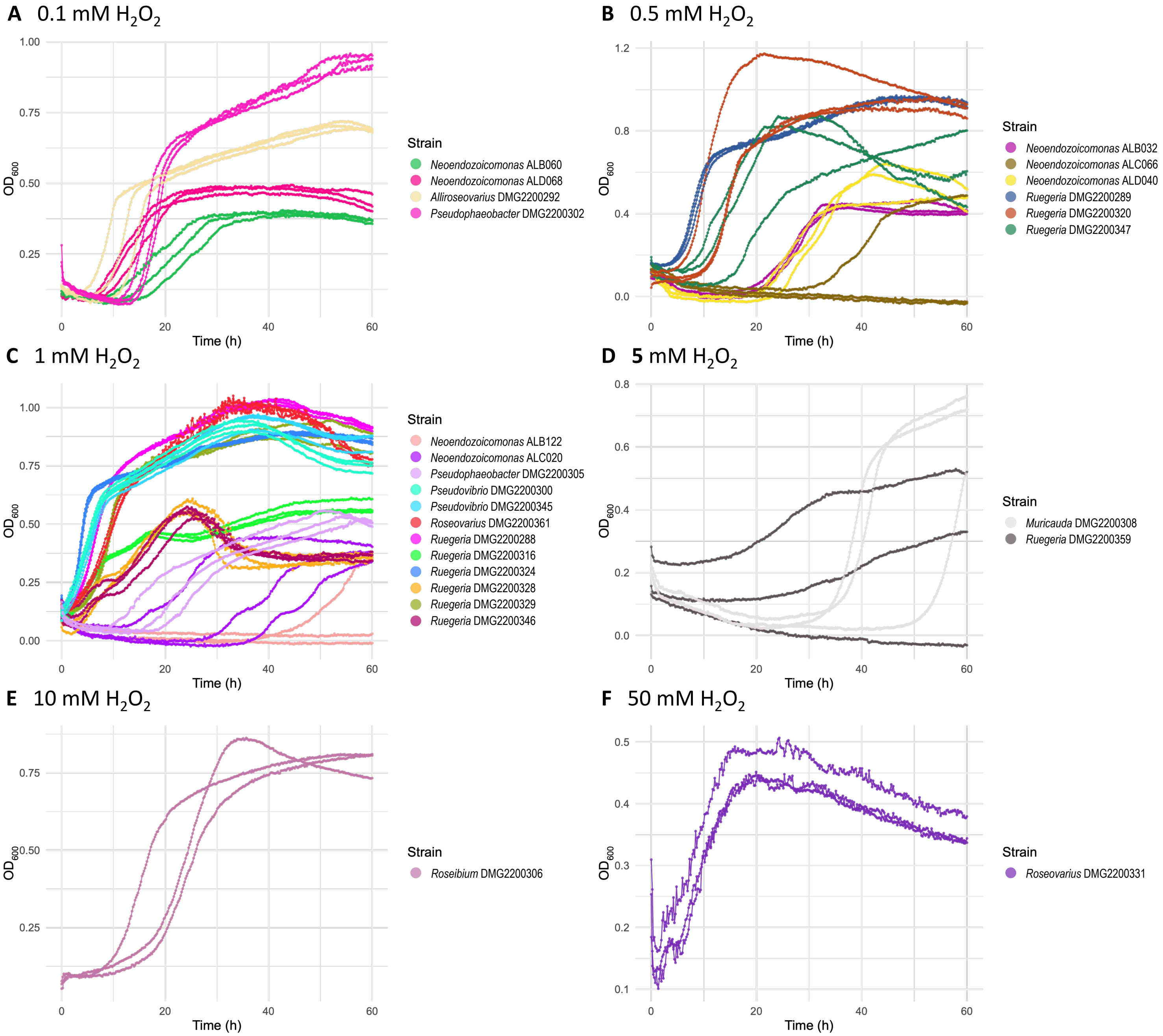


12

13 **Figure S3.** Growth curves of bacterial candidates at their maximum tolerated $H_2O_2$ concentration: (A) 0.1 mM, (B) 0.5 mM, (C) 1 mM, (D) 5 mM, (E) 10 mM, and (F) 50 mM $H_2O_2$.
